# FRET and LRET Biosensors for Cell-based Imaging and Screening of Rac1 Activation

**DOI:** 10.1101/2021.06.25.449998

**Authors:** Ha Pham, Mona Hoseini Soflaee, Andrei V Karginov, Lawrence W. Miller

## Abstract

Rac1 is a key regulator of several cell signaling pathways and dysregulated Rac1 activation has been implicated in cancer. Genetically encoded Förster resonance energy transfer (FRET) biosensors with enhanced dynamic range enabled live cell fluorescence imaging of Rac1 activity and a cell lysate-based assay of Rac1 inhibition in 96-well plates. We prepared HEK293 cell lines that stably expressed polypeptides with a general domain sequence (*N*- to *C*-terminus) of 1) FRET acceptor; 2) Rac/Cdc42 binding domain of human p21 protein kinase A (residues 68-150); 3) a linker domain; 4) FRET donor; and 5) full-length Rac1. Activated Rac1 binds to the protein kinase A domain, bringing donors and acceptors close together to increase FRET. We evaluated the effects on FRET signal dynamic range of alpha helical linkers comprised of alternating repeats of roughly four glutamate and four arginine or lysine residues. So-called ER/K linkers had limited effects on conventional FRET biosensors that incorporated the fluorescent protein (FP) pairs mCerulean/YPet, or mTFP1(cp227)/mVenus(cp229). Fluorometric measurements of cells that co-expresssed biosensors with positive (TIAM1) or negative (RhoGDI) Rac1 regulators revealed significant dynamic range enhancement in only one FP construct (mCerulean/YPet with 20 nm ER/K linker) relative to an analogous structure that incorporated an unstructured linker. We transfected this construct into a cell line that stably expressed a rapamycin-inducible c-Src analog (RapR-Src) and observed activation of Rac1 at protruding edges following rapamycin stimulation. In cells that expressed lanthanide-based FRET (LRET biosensors) that incorporated a luminescent terbium complex donor and GFP fluorescent acceptor, time-gated luminescence (TGL) measurements showed substantial gains in dynamic range that increased with linker length (up to 1200%). We robustly detected small molecule Rac1 inhibition following lysis of LRET biosensor-expressing cells grown directly in 96-well plates. The results herein highlight the potential of FRET and LRET biosensors with ER/K linkers for cell-based imaging and screening of protein activities.

## INTRODUCTION

Rac1 is a member of the Rho family of GTPases that regulate diverse cellular processes including actin remodeling, adherence junction formation, and establishment of cell polarity.^*1,2*^ Rac1 signaling also mediates transcriptional activation,^*3*^ cell cycle progression,^*4*^ G2/M checkpoint activation, and DNA damage response.^*5,6*^ The varied functions of Rac1 require a complex regulation of its subcellular location and interactions with multiple downstream targets.^*7*^ Rac1 is activated by the release of guanosine diphosphate (GDP) and the loading of guanosine triphosphate (GTP), which is catalyzed by upstream regulator guanine nucleotide exchange factors (GEFs).^*8*^ Besides GEFs, other major regulators which control Rac1 GTP/GDP binding status are GTPase activating proteins (GAPs) which accelerate the hydrolysis of GTP into GDP to turn Rac1 off ^*9*^ and guanine nucleotide dissociation inhibitors (GDIs) which bind to GDP-associated Rac1 and chaperone it in the cytoplasm in an inactive state.^*10*^

Fluorescence-based biosensors have played a vital role in Rac1 studies by enabling quantification of the spatiotemporal dynamics of Rac1 nucleotide-binding states in living cells. Most of these biosensors are recombinantly expressed fusion proteins that rely on the phenomenon of Förster resonance energy transfer (FRET) to read out changes in protein binding or conformation. FRET is non-radiative, dipole-dipole energy transfer that occurs when three conditions obtain: i) a luminescent donor molecule or nanoparticle resides within ~10 nm of an acceptor chromophore; ii) the donor’s fluorescence emission spectrum overlaps the acceptor’s absorbance spectrum; and iii) there is optimal alignment between the FRET partners’ respective emission and absorbance transition dipole moments^*11*^. FRET changes may be dynamically detected with spectrometers or appropriately configured microscopes as reductions in donor emission intensity or lifetime or increases in acceptor emission intensity coincident with donor excitation, if the acceptor species itself is fluorescent. In 2000, Kraynov et al. developed the first Rac1 biosensor, called FLAIR (fluorescence activation indicator for Rho proteins), to monitor the interaction between two fluorescently labeled proteins: i) a chimera of Rac1 fused to a fluorescent protein (FP); and ii) the p21-binding domain of Pak1 (PBD) labeled with the dye, Alexa546.^*12*^ Pak1 is a downstream effector that binds to the active state of Rac1. With the FLAIR construct, Rac1 activation increased the proportion of Rac1-GFP bound to Pak1-Alexa546 which in turn resulted in enhanced FRET between the GFP donor and Alexa546 acceptor.

Since the report of FLAIR, steady improvements have been made to Rac1 biosensors and the performance of FRET-based bioimaging more generally.^*13*^ Intensity-based FRET microscopy generally suffers from low signal-to-noise (S/N) and dynamic range because overlapped fluorophore spectra cause donor-sensitized, acceptor emission signals to be comingled with donor and directly excited acceptor fluorescence.^*11*^ The deleterious effects of spectral overlap worsen when imaging two freely diffusing fluorophores, especially when there is a large localized difference in either donor or acceptor abundance. Single-chain biosensor designs simplify FRET imaging and data analysis by incorporating sensing domains and fluorophores into a single polypeptide to maintain a 1:1 ratio of donors and acceptors.^*14*^ However, a single-chain affinity biosensor can fold into an off-state, or low-FRET, conformation that positions donor and acceptor in close proximity and yields a high baseline signal. Consequently, the dynamic range of many biosensors – i.e., the maximum observed difference between active and inactive conformations – rarely exceed 50%.^*15*^ A number of innovative single-chain Rac1 biosensors have been reported that incorporate optimized linker designs or circularly permuted fluorescent proteins (cpFPs) to enhance performance, with some reported examples exhibiting dynamic ranges of 150% or more.^*16-21*^

Here, we describe genetically encoded biosensors that enable single-cell imaging or higher throughput, cell-based assays of Rac1 activity in 96-well plates. Upon stable expression in HEK293T cells, each biosensor incorporates full-length Rac1, PDB and FRET partners into a single fusion protein. Rac1 activation causes PBD and GTP-bound, full-length Rac1 domains to bind one another, and the attendant conformational change repositions FRET partners to increase sensitized emission. Similar to our previously described designs,^*22, 23*^ these Rac1 biosensors incorporate two uncommon features that can enhance sensitivity and dynamic range. For one, alpha helical, ER/K linker segments separate affinity binding domains and FRET partners, maintain sensors in an extended conformation in the unbound (low activity) state, and thereby reduce baseline FRET signals. For another, *in situ* labeling of sensor peptides with luminescent Tb(III) complexes enables time-gated luminescence (TGL) detection of luminescence resonance energy transfer (LRET) between Tb(III) donors and GFP acceptors that is characterized by high signal-to-background (S/B).

We evaluated the effects of ER/K linker length (10, 20 or 30 nm) on the dynamic ranges of biosensors that featured three different donor/acceptor pairs: i) mCerulean/Ypet; ii) circularly permutated (cp) mTFP1/cp Venus; and iii) Tb(III)/EGFP). We co-expressed each biosensor configuration with regulatory proteins that either increased (TIAM1) or decreased (GDI) Rac1 activity and measured fluorescence spectra (for FP-based sensors) or TGL of Tb(III) and Tb(III)-to-GFP emission (for LRET sensors). Dynamic range was estimated as the difference between donor-ratioed, sensitized acceptor emission intensity seen in Rac1-active and inactive cell lines. For Tb(III)-based sensors, dynamic range increased with ER/K helix length to a maximum 1100% for sensors with a 30 nm linker. Large signal differences and high apparent Z’ factors were also observed in a 96-well plate, LRET assay that measured Rac1 inhibition in cell lysates. FP-based sensor performance in fluorometric assays did not correlate with ER/K linker length. However, ER/K-based, Ypet/mCerulean Rac1 biosensors exhibited substantial signal changes up to 125%. Using FRET microscopy and an engineered cell line that stably expressed a Ypet/mCerulean sensor with a 20 nm ER/K linker, we observed robust Rac1 activation near protruding edges of stimulated cells. These results, along with our earlier studies,^*22*^ demonstrate that FRET or LRET biosensors with ER/K linkers are a robust platform for engineering sensitive single cell imaging studies or higher-throughput cell-based assays of protein function in live cells.

## RESULTS AND DISCUSSION

### Biosensor Design

Our sensor design relies on the interaction between the Rac/Cdc42 binding domain of human p21 protein kinase A (residues 68-150; PBD) and the active, GTP-bound form of Rac1 to report changes in Rac1 activity. Each sensor fusion protein incorporated five domains, ordered from *N*- to *C*-terminus as follows: i) a FRET acceptor FP; ii) PBD; iii) an ER/K linker sequence (length of 10, 20 or 30 nm); iv) either a FRET donor FP or eDHFR that binds to a trimethoprim (TMP)-Tb(III) complex conjugate; and v) full-length Rac1 (**Figure 1A**). We placed Rac1 at the *C*-terminus to retain the CAAX box that localizes native Rac1 at the plasma membrane and facilitates regulation by RhoGDI.^*12*^ We prepared biosensors with three different FRET/LRET pairs that included mCerulean/Ypet, cp mTFP1/cp Venus, and eDHFR(Tb)/EGFP (nine total sensors, **Figure 1B**). We chose mCerulean/YPet and mTFP1 (cp227)/Venus (cp229) as FP FRET partners because the former has been shown to exhibit high dynamic range, and the latter was previously used in the Rac1-2G biosensor,^*16*^ which had the best spectral property among Rac1 biosensors made with cp variants of mTFP1 and Venus.

**Figure 1.**
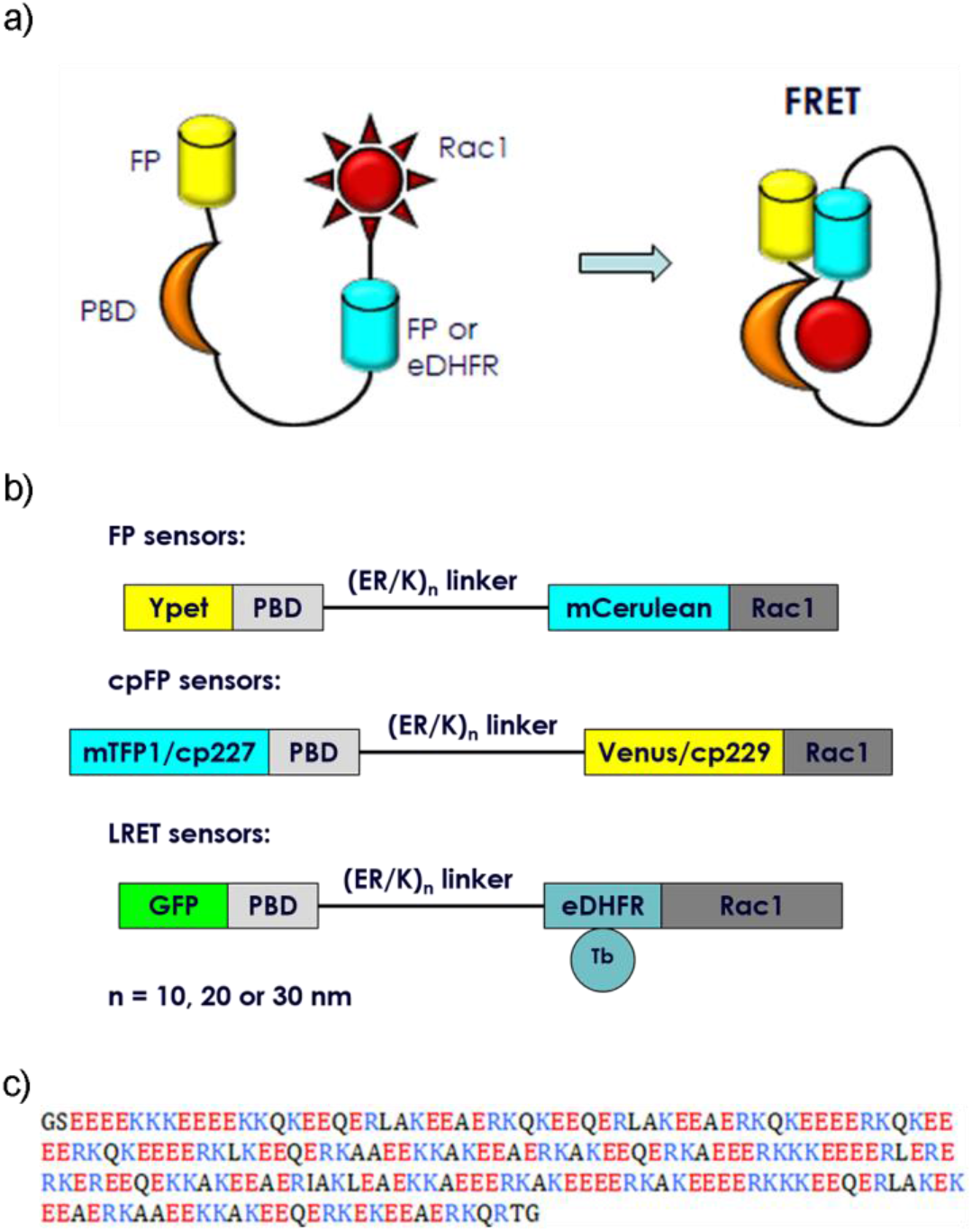
Designation of FRET and LRET Rac1 biosensors. (A) Schematic representation and (B) mammalian expression constructs of single-chain FRET or LRET Rac1 biosensors. A series of nine sensors was constructed in which alpha-helical ER/K linkers of different length (10 nm, 20 nm and 30 nm) were combined with three fluorophore pairs: mCerulean/Ypet, circularly permutated mutant (cp227) of mTFP1/circularly permutated mutant (cp229) of Venus, and Tb(III) complex/EGFP. (C) Sequence of a 30 nm alpha-helical ER/K linker that includes alternated repeats of approximately four negatively charged glutamates (red) and four negatively charged arginines or lysines (blue). The linker is rigid to keep protein components far apart in OFF state. Stochastic breaks allow protein interaction in the biosensor ON-state and promote a large change in donor-sensitized acceptor emission.

We previously showed that biosensors which incorporate Tb(III) complexes as LRET donors, GFP as LRET acceptors and ER/K linkers can be used to craft cell-based TGL assays of protein-protein interactions that exhibit exceptional sensitivity and dynamic range.^*22*^ Inclusion of an eDHFR domain permits self-assembly with TMP-Tb(III) complex conjugates inside live cells or in cell lysates. TGL detection of LRET offers high signal-to-background ratios that derive from the ms-scale emission decay times of Tb(III) luminescence and Tb(III)-to-GFP sensitized emission. TGL plate readers or microscopes implement a brief delay (~μs) between pulsed, near-UV excitation and detection that nearly eliminates non-specific fluorescence (with ~ns decay times) from cells, library compounds or directly excited acceptors.

The ER/K linker forms an extended alpha helix that keeps apart FRET partners and binding domains to minimize baseline energy transfer when the sensor is in the unbound or off state. It is presumed that random breaks in the helix permit close approach and binding of terminal affinity domains, and the proportion of biosensors in the closed and open conformations depends solely on the ER/K linker length and the inherent affinity of the binding partners.^*24*^ We investigated the effects of ER/K linker length on dynamic range by preparing biosensors with linkers of approximately 10 nm, 20 nm, or 30 nm (**Figure 1C**).

### Biosensor Characterization

We evaluated dynamic range by co-expressing all nine biosensors along with Rac1 upstream regulators and using a fluorometer (for FP or cpFP biosensors) or a TGL plate reader (for LRET biosensors) to read out emission signals. We transformed HEK293T cells to stably express biosensors under control of a tet-inducible promoter. The stably transformed cell lines were transiently transfected with plasmid DNA that encoded either Tiam1 or GDI to maintain sensors in the Rac1-active or inactive states, respectively. Following transient transfection and induction of biosensor expression in FP or cpFP cell lines, we obtained scanning fluorescence emission spectra of suspended cells. For plate reader assay, 293T cells co-expressing LRET biosensors and Tiam1 or GDI were seeded in a 96 well plate. A lysis solution containing a TMP-Tb(III) conjugate (TMP-TTHA-cs124) was added to the cells approximately ten min before TRL measurement of Tb(III) and Tb(III)-to-GFP emission intensity. We calculated biosensor dynamic range as (R_a_-R_i_)/R_i_, or ΔR/R_i_, where R_a_ and R_i_ were the sensitized acceptor (FRET)-to-donor emission ratios obtained from spectra of cells expressing activated (with TIAM1) and inactivated (with GDI) biosensors, respectively. For comparison, we also analyzed FP-based Rac1 biosensors with flexible linkers or ER/K linkers (10, 20, 30 nm) including four that included mCerulian/YPet as the donor/acceptor pair as well as analogs of the previously reported Rac1-2G biosensor, comprised of (*N* to *C*), mTFP1(cp227), PBD, a linker (flexible, or ER/K), Venus(cp229), and Rac1 (**Figure 1B**).^*16*^

Based on our prior studies, we expected to see a large difference in donor-denominated FRET or LRET ratios between GDI and TIAM1 cells that increased with ER/K linker length. However, the incorporation of ER/K helical linkers had little effect on the performance of FP (mCerulean/YPet) or cpFP (mTFP1cp227/mVenuscp229) biosensors. The FRET ratios measured in mCerulean/YPet biosensor cells that co-expressed TIAM1 (On-state, Rac1 active) exceeded those measured in GDI co-expressing cells (Off-state, Rac1-inactive) by 96% (± 16%, mean ± sem), 125(±8)%, and 117(±17)% when ER/K linkers measured 10, 20 or 30 nm, respectively (**Figure 2A**). Only the mCerulean/YPet biosensor with a 20 nm ER/K linker showed a significantly (p < 0.01) improved dynamic range in comparison to the same configuration that incorporated an unstructured linker. in a statistically significant manner (128±8% vs. 97±7%). The dynamic range value that we observed for Rac1-2G (67%) was nearly same as the literature value (69%),^*16*^ and values for cpFP anlogs with ER/K linkers ranged from 52% to 73% with no correlation to linker length (**Figure** 2A).

**Figure 2.**
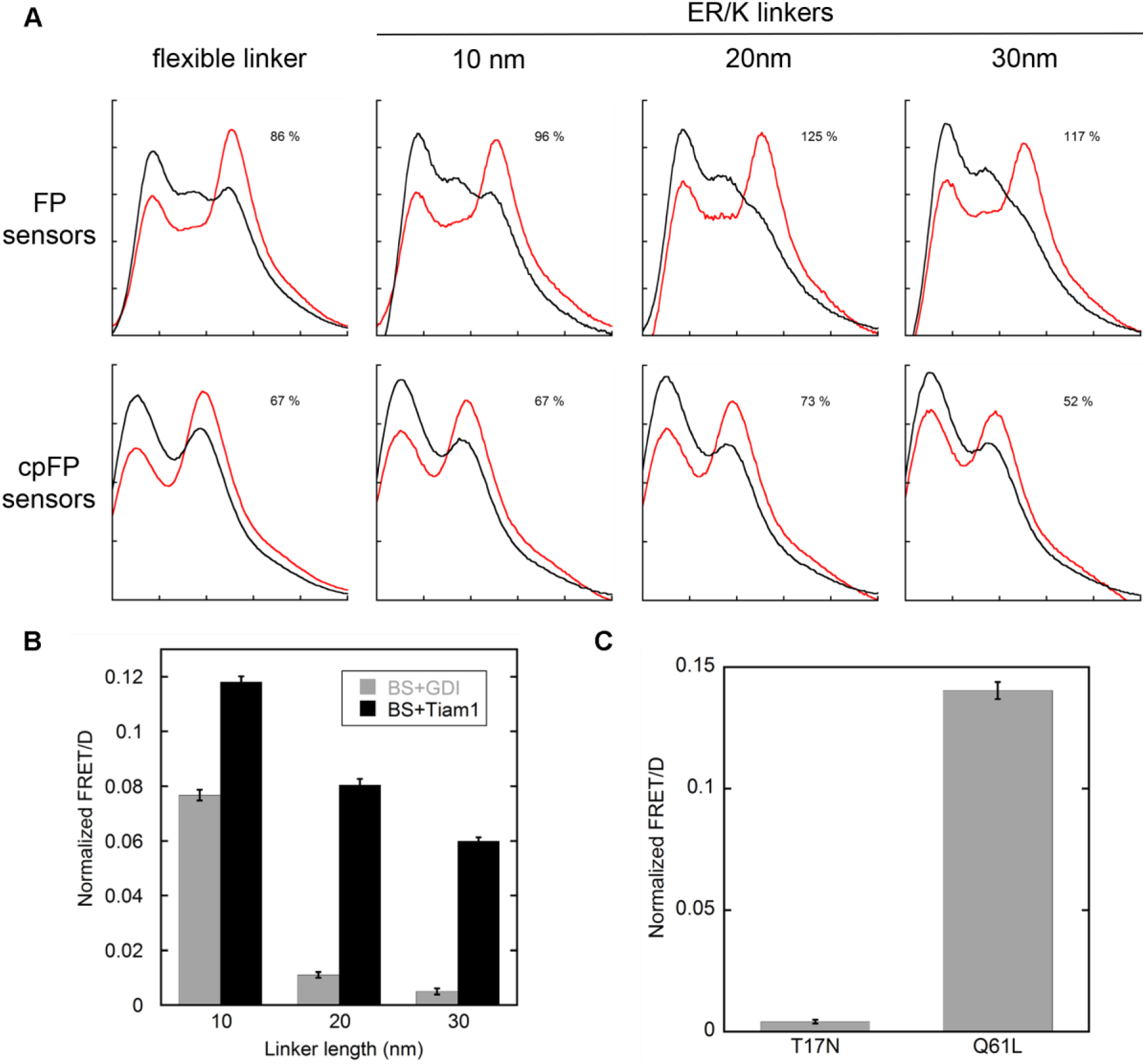
Evaluation of Rac1 biosensors by fluorometry and TGL luminescence. Rac1 biosensors were evaluated by co-expressing each sensor with Tiam1 (On state, red trace in A or black bar in B) or RhoGDI (Off state, black trace in A or gray bar in B) in 293T cells. (A) Cells co-expressing FP or cpFP sensors and the regulators were scanned for emission profiles in cell suspension using a cuvette-based fluorometer. ΔR/R_i_ values representing differences in FRET efficiency between the On and Off states are shown for the indicated biosensor. (B) Cells co-expressing LRET biosensors and the regulators were grown in a 96-well plate. Following overnight incubation, cells were treated with lysis buffer containing TMP-TTHA-cs124 (25 nM). The time-gated emission (gate delay, 0.2 ms) at 520 nm (Tb-to-GFP LRET) and 620 nm (Tb only) were measured using a time-resolved fluorescence plate reader. Substantially larger 520 nm/620 nm (FRET/D) ratios were observed in the positive control wells (Tiam1) relative to those seen in the negative controls (RhoGDI). (C) 293T cells expressing inactive (Rac1 T17N) or active (Rac1 Q61L) mutants of the LRET biosensor with 30 nm ER/K linker were grown in a 96-well plate and underwent the same lysis-buffer treatment and plate-reader measurement as in (B). Fluorescence spectra in (A) are representative of cells transiently expressing biosensors with transfection efficiency > 70%. Dynamic range values for cpFP sensors are from single experiments (transfections). For FP sensors, dynamic range values represent mean (n ? 3 transfections) Error values (sem) of FP sensor dynamic range: flexible linker, 7%; ER/K 10 nm, 16%; ER/K 20 nm, 8%; ER/K 30 nm, 17%. Bar graphs in (B), (C), sem (n = 16 wells).

By contrast, the observed dynamic ranges of Tb(III)-based biosensors strongly depended on ER/K linker length. LRET sensors with 10 nm, 20 nm, or 30 nm linkers exhibited maximum differences in ΔR/R_i_ values of 50%, 600%, and 1100%, respectively (**Figure 2B**). To further validate this result, we performed the same plate reader assay on cells that expressed active (Rac1 T17N) or inactive (Rac1 Q61L) mutants of the LRET biosensor with 30 nm linker. The difference in LRET/donor emission ratio between active and inactive mutants (1200 %) was similar to that seen in the Tiam1/GDI assay (**Figure 2C**).

The unique properties of the Tb(III) complex as a donor can partly explain the robust dynamic range of LRET sensors. In essence, the narrow emission bands of the Tb complex minimize bleedthrough, and the TGL detection of long-lived donor and FRET signals eliminates background fluorescence. Therefore, even without purifying proteins after lysing cells, our plate reader assay still gave distinct signals between two states of the biosensors. Moreover, the low protein concentration in a plate well (< 10 nM) is typically below the detection limit of FP or cpFP biosensors, highlighting another advantage of LRET-based sensors and TGL analysis.

The characterization data illustrate that longer linkers significantly decreased background, or Off-state LRET while less dramatically reducing On-state LRET (**Figure 2B**). The effect of ER/K linkers on the sensor properties can be understood by considering their conformational behavior. Unlike unstructured, flexible middle linkers, alpha-helical ER/K linkers predominantly adopt an extended conformation in the unbound state. The presumably rigid structure can explain why Off-state LRET diminishes with linker length. However, increased linker length also reduces the effective concentration of the binding partners located at either end,^*24*^ which may explain why maximal (On-state) LRET signals also decrease with linker length in the Tb(III)-based biosensors.

While ER/K linker length substantially impacted the performance of LRET Rac1 biosensors, they had minimal effect on that of FP- or cpFP-based biosensors. The strong influence in LRET biosensors may be due to the long distance of LRET, which causes high baseline signals and requires a long rigid linker for extended conformation in the Off state. Second, orientation factors are essential in FRET efficiency between FPs, especially cpFPs. In fact, the purpose of using cpFPs is to alter the arrangement of protein elements in the sensors so as to produce favorable spatial orientation (κ^2^). Thus, the rigidity of ER/K linkers may counteract this purpose by restricting the mobility of cpFP dipole toward unfavorable (κ^2^ < 2/3) orientation. In general, a flexible linker is preferred in FP and cpFP sensors because it provides little, if any, restriction on the tumbling and twisting of the fluorophores. In contrast, LRET does not depend on the dipole orientation of the donor or acceptor.^*25*^

### Biosensors in Live Cells

We next evaluated the performance of FP-based sensors for FRET microscopic imaging of Rac1 in HeLa cells. In imaging studies, Rac1 activity in sensor-expressing cells has been induced with growth factors (EGF, PDGF) or other extracellular stimuli. However, this approach triggers a multitude of parallel signaling pathways and often a brief, transient activation of Rac1 which significantly complicates the analysis of sensor activity. To simplify our imaging characterization experiments, we applied a recently developed protein engineering method that employed a rapamycin-regulated allosteric switch to regulate tyrosine kinase c-Src (Src) activation.^*26*^ In this way, activation of Rac1 could be controlled indirectly by rapamycin-induced Src (RapR-Src) activation. For live-cell imaging, cells co-expressing the mCerulean/YPet Rac1 biosensor with 20 nm ER/K linker and RapR-Src constructs were seeded on a fibronectin-coated chamber slide. Cells were imaged every 2 min for 20 min before and 40 min after adding rapamycin. We observed robust Rac1 activation near protruding edges of stimulated cells following rapamycin addition. Cells started vigorous spreading and protrusion a few minutes after stimulation. Increased sensor activity was consistently observed wherever protrusions were extending (**Figure 3**).

**Figure 3.**
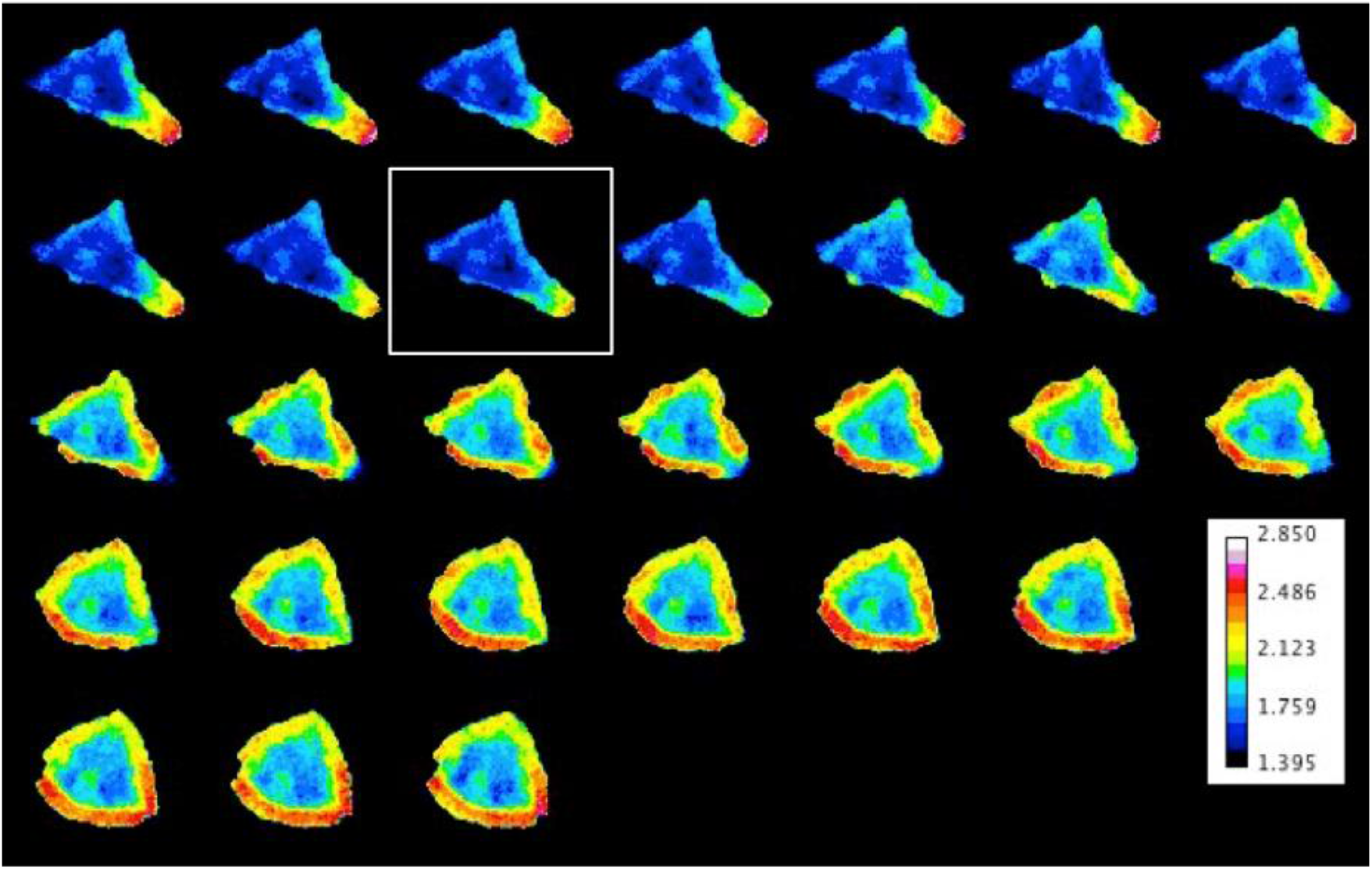
Live-cell imaging of Rac1 biosensors. A HeLa cell co-expressing the FP Rac1 sensor with the 20 nm ER/K linker (transient transfection) and RapR-Src constructs (stably transfection) was imaged for every 2 min in 1 hour. Rapamycin (500 nM) was added after 20 min (indicated by the square). The montage shows FRET/mCerulean ratio images of the biosensor in a living cell. Warmer colors reflect higher local activity of Rac1.

### Detection of Rac1 inhibition in multi-well plates

Increasing evidence supports the involvement of Rac1 signaling alterations in cancers.^*27, 28*^ Because mutations in Rac1 proteins are sporadic, the mechanism of Rac1 in cancer likely occurs through its overexpression or hyperactivity.^*29*^ In fact, overexpression of Rac1 and its hyper-activation caused by overexpression of Rac1 specific GEFs such as Tiam1 and Vav1 have been observed in various tumor types, including pancreatic cancer.^*30, 31*^ Accordingly, the inhibition of Rac1 is speculated to have an antiproliferative effect on cancer cells.^*28*^ Several small molecules have been reported as Rac1 inhibitors.^*32–36*^ NSC23766 and EHT 1864 were among the first developed Rac1 inhibitors with the capability to discern from other Rho family GTPases, such as Cdc42 or RhoA.^*35 36*^ The former impedes Rac1 activation by occupying the binding location of two Rac1 GEFs: Trio and Tiam1,^*35*^ while the latter keeps Rac1 in an inactive state and prevents its binding to downstream effector.^*36*^ At this point in time, small molecule inhibitors that target Rac1 have been identified through structure-based virtual high throughput screening,^*32, 34*^ followed by pull-down assays to examine hits.^*37, 38*^ Although this method is realizable, the technical complication, high cost, and the missing of native environment during screening are among its disadvantages.

We sought to assess the potential of our LRET Rac1 biosensor for the detection and quantification of Rac1 inhibition in a multi-well plate format. We measured the effects of NSC23766 and EHT 1864 as inhibitors of Rac1 activation. 293T cells were seeded into 96-well plates (9000 cells/well), followed by transient transfection of the biosensor and Tiam1 on the next day. Tiam1 was overexpressed in cells to activate Rac1, mimicking the deregulation of this regulator in cancer cells.^*39*^ 48h after transfection, cells were incubated with growth media containing inhibitor (either NSC23766 and EHT 1864, 50 uM) for four hours or overnight. Subsequently, growth media was exchanged for a lysis buffer that contained TMP-Lumi4-Tb (25 nM), and the plate was incubated at room temperature for 10 min. Negative control wells went through the same treatment but without addition of inhibitor. Following incubation, the time-gated Tb(III)-to-GFP and Tb(III) emission signals were measured at 520 nm and 615 nm, respectively. The 520 nm signal from each well was divided by the 615 nm signal to minimize well-to-well variability resulting from differences in probe amounts or sample absorbance. Then, the mean 520/615 emission ratio from 12 control wells containing non-expressing cells and lysis buffer solution (25 nM TMP-Lumi4-Tb, no rapamycin) was subtracted from each sample well to yield a background-corrected, LRET/Tb ratio.

The LRET Rac1 biosensor with 30 nm ER/K linker responded strongly to Rac1 inhibitors in the 96-well plate assay. After four-hour incubation, full inhibition with 50 uM EHT 1864 yielded LRET/Tb signal decrease of more than 70%. The ratio went down to the same level as the biosensor without Tiam1 activation (**Figure 4A**). Compared to EHT1864, NSC23766 partially inhibited the Rac1 activity after an overnight incubation (**Figure 4B**). We calculated apparent Z’ factors to assess statistical robustness and found that inhibition assays with overnight incubation of EHT 1864 or NSC23766 exhibited Z’ factor values of 0.69 and 0.19, respectively.

**Figure 4.**
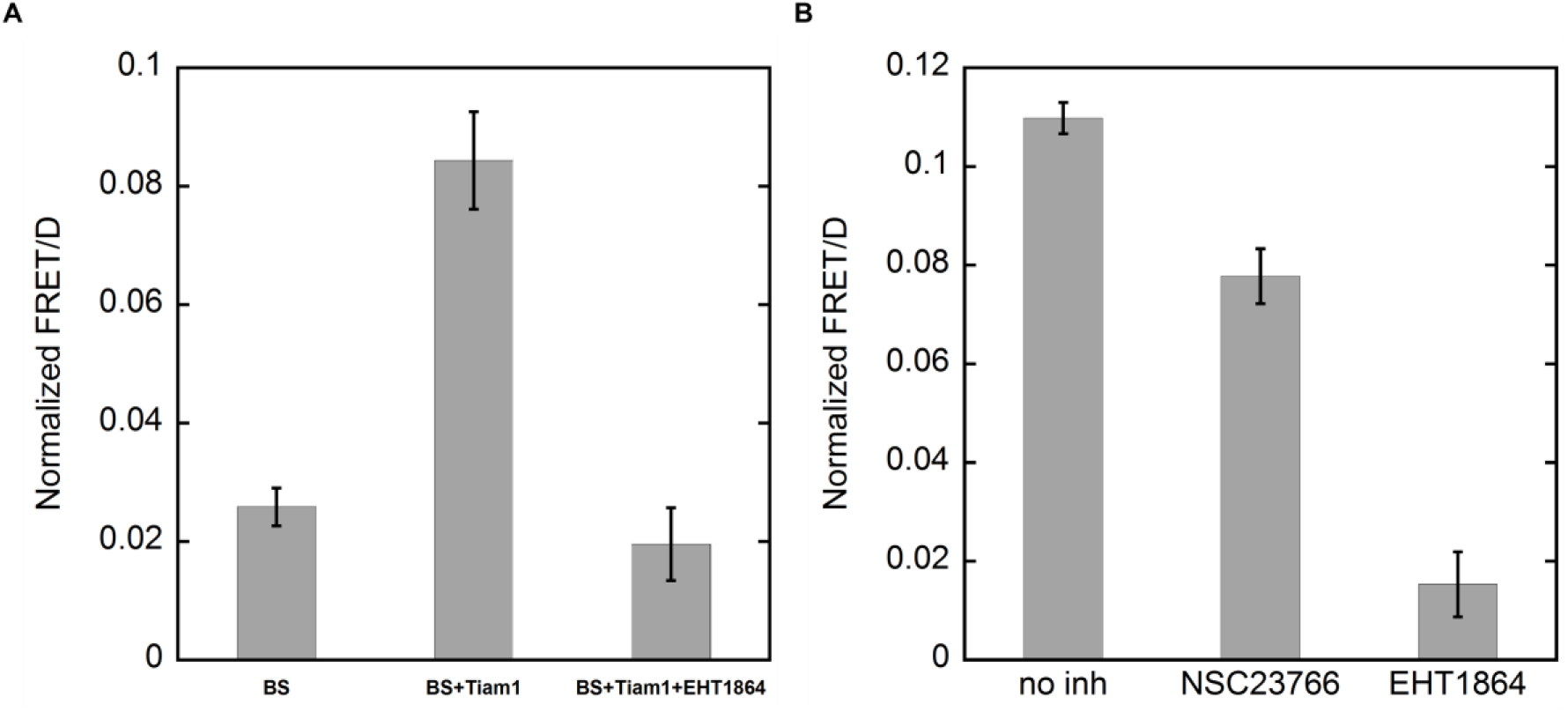
Large reductions were observed in multi-well plate when NSC23766 and EHT 1864 inhibit Rac1 activation. A. 293T cells expressing an LRET Rac1 biosensor (30 nm ER/K linker) alone (BS) or with Tiam1 (BS+Tiam1) were grown in 96-well. Time-gated measurements were obtained following 4h incubation with EHT 1864 and addition of a lysis buffer containing TMP-TTHA-cs124. B. 293T cells co-expressing the sensor and Tiam1 were grown in 96-well overnight with or without inhibitors in the media. Time-gated measurements were taken after adding the lysis buffer as in A. Bar graphs depict mean FRET/donor (D) ratio measured for each condition (n=16). Error bars, sem.

## CONCLUSION

LRET biosensors’ extraordinary dynamic range stems from time-gated detection of LRET that eliminates non-specific fluorescent background and from the incorporation of ER/K linkers that maintains the donor and acceptor far apart in the open sensor configuration. These features enable robust detection of Rac1 activation or inhibition in cell lysates in 96-well plates and support medium or high throughput detection of small molecules inhibitor for Rac1 or any other proteins, especially proteins that are difficult to purify or isolate from intact environment. Moreover, Tb(III) can sensitize differently colored acceptors, offering the potential for multiplexed imaging or analysis. While the dynamic range of LRET-based, affinity biosensors is clearly affected by the presence and length of ER/K linkers, the extent to which inclusion of a helical linker benefits conventional FRET biosensors is less clear. Taken together, the results presented here show that the LRET biosensor toolkit is versatile when used to improve existing biosensor templates.

## EXPERIMENTAL METHODS

### Materials

293T cells (CRL-3216), HeLa cells (CCL-2), and MEF cells (SCRC-1040) were from ATCC. Dulbecco’s modified eagle medium with 4.5g/L glucose (DMEM, 10-013CV), Dulbecco’s phosphate buffer saline (DPBS, 21-030 and 21-031), and 0.05% trypsin/2.21 mM EDTA (Corning, 25-053-Cl) were purchased from Corning cellgro ®. DMEM (without phenol red, 21063), HEPES (15630-080) and Lipofectamine 2000 (11668-027) were purchased from Invitrogen^TM^. FBS (S11150) was purchased from Atlanta Biologicals. Hygromycin (sc-29067) was purchased from Santa Cruz Biotechnology. BSA (700-107P) was purchased from Gemini Bio-products. Rapamycin (553211-500UG) and Fibronectin bovine plasma (F1141) were purchased from Millipore. NADPH (N0411) and doxycycline (D9891) were purchased from Sigma. DMSO (D128-500) was purchased from Fisher Chemical. Patent V blue sodium salt (21605) was purchased from Fluka. In-Fusion HD cloning kit (638909) was purchased from Takara. All enzymes and buffers used in cloning were purchased from New England Biolabs. pUSE-RapR-Src-as2-mCherry-myc and pUSE-ipep-FRB* plasmids were a gift from Dr. Andrei Karginov’s lab. FP biosensor with flexible linker was a gift from Dr. Hahn’s lab at the University of Noth Carolina at Chapel Hill. As for luminescent Tb(III) complexes. Heterodimers of trimethoprim linked to luminescent Tb(III) complexes (TMP-cs124-TTHA,^*40, 41*^ and TMP-Lumi4^*40*^) and a cell permeable variant conjugated to oligoarginine (TMP-Lumi4-R9)^*42*^ were prepared as previously reported.

### Cell culture

HeLa cells were maintained in DMEM (1.0 g/L glucose) supplemented with 10% FBS, 1X MEM non-essential amino acids and 15 mM HEPES at 37 °C and 5% CO2. The cells were passaged with 0.25% trypsin/2.21 mM EDTA. 293T cells were maintained in DMEM (4.5 g/L glucose) supplemented with 10% heat-inactivated FBS and 2mM L-glutamine at 37 °C and 5% CO2. The cells were passaged with 0.05% trypsin/2.21 mM EDTA. NIH 3T3 cells were maintained in DMEM (4.5 g/L glucose) supplemented with 10% FBS at 37 °C and 5% CO2. The cells were passaged with 0.05% trypsin/2.21 mM EDTA.

### Plasmids

All DNA constructs were sequenced by the UIC Research Resources Center (RRC).

**pTriEX-Ypet-PBD-(ER/K)_n_-mCerulean-Rac1 (n = 10nm, 20nm, or 30 nm)** was prepared by subcloning from pTriEX-Ypet-PBD-mCerulean-Rac1. The source vector was provided by Hahn lab at the University of North Carolina at Chapel Hill. HindIII to NotI fragments encoding 10 nm, 20 nm, or 30 nm ER/K linker were amplified using the following primer pairs respectively: 5’-ACT GAA GCT TCA GGA AGC GGA GAA GAG GAA GAG A-3’ and 5’-TGA TGC GGC CGC CAG AGC CCT TCT TCT TGC G-3’; 5’-ACT GAA GCT TCC GGA GGA TCC GAA GAG GAG GA-3’ and 5’-TAA TGC GGC CGC CAG AGC CAC CGG TCT CT-3’; 5’-AGC AAA GCT TCT GGA TCC GAA GAG GAG GAG A-3’ and 5’-CTT AGC GGC CGC CAC CGG TTC TCT GTT TTC GC-3’. The linker fragment was then inserted between the HindIII site and the NotI site in the source vector.

**pTriEX4-mTFP1/cp227-PBD-(ER/K)_n_-Venus/cp229-Rac1 (n = 10nm, 20nm, or 30 nm)** was subcloned from pTriEX4-mTFP1/cp227-PBD-Venus/cp229-Rac1. The source vector was purchased from Addgene. XmaI to NotI fragments encoding 10 nm, 20 nm or 30 nm ER/K linker were amplified using the following primer pairs respectively: 5’-TAA TCC CGG GGG AAG CGG AGA AGA GGA AG-3’ and 5’-TAA TGC GGC CGC GCC AGA GCC CTT CTT CTT GC-3’; 5’-TAA TCC CGG GGG AGG ATC CGA AGA GGA GGA GAA-3’ and 5’-TAA TGC GGC CGC AGA GCC ACC GGT CTC TTC-3’; 5’-CGT ACC CGG GGG AGG ATC CGA AGA GGA GGA-3’ and 5’-CTT AGC GGC CGC ACC GGT TCT CTG TTT TCG-3’. Those fragments were then inserted between the XmaI site and the NotI site in the source vector.

**pTriEX4-EGFP-PBD-(ER/K)_n_-eDHFR-Rac1 (n = 10nm, 20nm, or 30 nm)** was subcloned by replacing mTFP1/cp227 and Venus/cp229 in pTriEX4-mTFP1/cp227-PBD--(ER/K)n-Venus/cp229-Rac1 with EGFP and eDHFR. The EGFP fragment was subcloned between NcoI and BspEI sites using the primer pair: 5’-TCG CCA CCA TGG TGA GCA AG-3’ and 5’-TTC GAA GCT TGA GCT CGA GAT CTG-3’. The eDHFR fragment was subcloned between NotI and KpnI sites using the primer pair: 5’-ATG CAG CGG CCG CCA TGA TCA GTC TGA TTG CGG CGT TA-3’ and 5’-ATC GAG GTA CCA GAC CGC CGC TCC AGA ATC TCA-3’.

### Doxycycline-inducible constructs

To generate **pPBH-TRE_tight_-mTFP1/cp227-PBD-(ER/K)_20_-Venus/cp229-Rac1**, the entire biosensor cassette was subcloned to PiggyBac Transposon vector system (SBI System Biosciences) by In-Fusion HD cloning kit. The biosensor construct was amplified using the primer pair: 5’-ACT CTG CAG TCG ACG GTA CCT TAT TTA CAA TCA AAG GAG ATA TAC CAT GGT GAG C-3’ and 5’-AGC TTA TCG ATG CGG CCG CGG ATA GGC AGC CTG CAC CTG A-3’. To generate **pPBH-TREtight-EGFP-PBD-(ER/K)_30_-eDHFR-Rac1**, the gene encoding PBD-30nm ER/K linker, and eDHFR-Rac1 were subcloned with infusion primers to a PiggyBac vector containing EGFP. A 918 bp fragment encoding PBD-30nm ER/K linker was amplified by PCR using the infusion primers 5’-ACT CTG CAG TCG ACG GTA CCT CCG GAC TCA GAT CTG AGC TC-3’ and 5’-CTG ATC ATG CCA GAA CCG GTT CTC TGT TTT CGC TCT-3’. A 1071 bp fragment encoding eDHFR-Rac1 was amplified by PCR using the infusion primers 5’-TTC TGG CAT GAT CAG TCT GAT TGC GGC G-3’ and 5’-ATG CGG CCG CGC TAG CCT ACA ACA GCA GGC ATT TTC TCT TCC-3’. These two fragments were inserted between the KpnI site and the NheI site the destination vector by infusion enzyme. Plasmid integrity was confirmed by direct sequencing. To generate the dual-promoter plasmid: **pPBH-TRE_tight_-RapR-Src-as2-mCherry-myc and pCMV-ipep-FRB\***, the gene encoding RapR-Src-as2-mCherry-myc and (CMV Promoter)-ipep-FRB*-(bGH Poly(A) Signal Sequence) from pUSE-RapR-Src-as2-mCherry-myc and pUSE-ipep-FRB*, respectively, were subcloned to a PiggyBac vector. The fragment encoding RapR-Src-as2-mCherry-myc was amplified by PCR from pUSE-RapR-Src-as2-mCherry-myc by using the primer 5’ – ATG CAG CTA GCA TCA TGG GCA GCA ACA AGA GC – 3’ (BmtI coding strand) and 5’ – ATG CAT CTA GAA TCA CCA GTT TCT TCC GGA CTT GTA C – 3’ (XbaI non-coding strand). This fragment was inserted at the BmtI and XbaI site in the PiggyBac vector. The fragment encoding ipep-FRB* was amplified by PCR from pUSE-ipep-FRB* by using the primer 5’ - GCG GCG CCC TGC CCG TCC CAC CAG GTG AGT TCC GCG TTA CAT AAC TTA CGG – 3’ (SexAI, coding strand), and 5’ – GGC CGG TTA CCG CCT GTT GAC CTG GTC GCG TTA AGA TAC ATT GAT GAG TTT GGA C – 3’ (SexAI, non-coding strand). This fragment was inserted at the SexAI site in pPBH-TREtight-RapR-Src-as2-mCherry-myc using In-Fusion® Cloning Kit.

### Fluorometry assay

pTriEX-Ypet-PBD-(ER/K)_n_-mCerulean-Rac1 and pTriEX4-mTFP1/cp227-PBD-(ER/K)n-Venus/cp229-Rac1 (n = 10nm, 20nm, or 30 nm) biosensor constructs were co-transfected with upstream regulators (Tiam1 or GDI) into 293T cells plated 1×10^5^ per well overnight in 6-well plates, using Lipofectamine 3000 (Invitrogen) according to the manufacture’s protocols. The total amount of DNA was 750 ng per well (150 ng of the biosensor and 600 ng of the regulator). At 48h after transfection, cells were detached with brief trypsin treatment and were resuspended in 500 μL of ice-cold PBS. Cell suspensions were transferred into a quartz cuvette. Their fluorescent spectrum was then recorded by a fluorometer. The FP biosensors and cpFP biosensors were excited at 433 nm and 460 nm respectively. Their emission spectra were recorded from 450 to 600 nm and 480 to 600 nm respectively. Spectra were background-subtracted with spectra of non-transfected cells and normalized according to their spectrum integral.

### Multi-well plate assays with permeabilized mammalian cells

To validate the LRET biosensors, pTriEX4-EGFP-PBD-(ER/K)n-eDHFR-Rac1 (n = 10nm, 20nm, or 30 nm) biosensor constructs were co-transfected with upstream regulators (Tiam1 or GDI) into 293T cells using Lipofectamine 2000. One day prior to transfection, cells were seeded at 9000 cells per well in a poly-L-lysine coated 96-well plate. The amount of DNA per well was 4.25 ng of the biosensor and 17 ng of the regulator. After 48h, growth media in the wells were discarded carefully and 70 uL of lysis buffer with TMP-cs124-TTHA (25 nM) was added into the wells. The plate was kept at room temperature in dark for 10 min prior to the first measurement. Background control wells received the same transfection media but no biosensor construct and the same lysis buffer with Tb(III) complex. The emission signals were background-subtracted and FRET/acceptor and FRET/ donor emission ratios were calculated.

### Inhibition assay with permeabilized mammalian cells

pTriEX4-EGFP-PBD-(ER/K)_30_-eDHFR-Rac1 biosensor construct were co-transfected with upstream regulators (Tiam1 or GDI) into 293T cells using Lipofectamine 2000. One day prior to transfection, cells were seeded at 9000 cells per well in a poly-L-lysine coated 96-well plate. The amount of DNA per well was 4.25 ng of the biosensor and 17 ng of the regulator. After 24h, cells were incubated with 100 uL of growth media containing NSC23766 or EHT1864 inhibitor (final concentration 50 uM) for 4 hours or overnight. Negative control wells received growth media without the inhibitor. Next, growth media in the wells were discarded carefully and 70 uL of lysis buffer with TMP-cs124-TTHA (25 nM) was added into the wells. The plates were kept at room temperature in dark for 10 min prior to the first measurement. Background control wells received the same transfection media but no biosensor construct and were treated the same as sample wells for the rest of the experiment. The emission signals were background-subtracted and FRET/acceptor and FRET/ donor emission ratios were calculated.

### Stable expression of biosensor plasmids

Rac1 biosensor constructs in PiggyBac vector system were transfected into MEF cells using Lipofectamine 2000. Hygromycin selection was applied to generate the stable cell lines. The cells were then FACS sorted to obtain a population of uniform but medium biosensor expression levels. For imaging experiments, cells were induced by adding 100 ng/mL doxycycline and appropriate biosensor expression levels were achieved at 24h following the induction.

### Cell imaging of cpFP Rac1 biosensor

HeLa cells stably expressing RapR-Src-as2-mcherry and ipep-FRB were plated on a fibronectin-coated 25 mm–diameter glass coverslip by placing the coverslip inside a well of a 6-well plate and adding 35,000 cells/well. Cells were incubated for 2–4 h. Before imaging, coverslips were placed into an Attufluor Cell Chamber (Invitrogen, catalog no. A78-16) with Ham’s f12 K medium (no red) containing 10mM HEPES, DL lactate, and Oxy Fluor, supplemented with 1% (vol/vol) FBS. Live-cell imaging was done using an Olympus IX-83 microscope controlled by Metamorph software and equipped with a heated stage (Warner Instruments), Olympus UPlanSAPO 40× (oil, N.A. 1.25) objective, Xcite 120 LED (Lumen Dynamics) light source, and Image EMX2 CCD (Hamamatsu) camera. We thank Dr. Andrei Karginov at Department of Pharmacology, UIC for helping and providing the equipment for this experiment.

### Cell imaging of LRET Rac1 biosensor

Time-gated luminescence images were acquired using a previously described epi-fluorescence microscope (Axiovert 200, Carl Zeiss, Inc.).^*43, 44*^ For each time-gated image acquisition, the signal from multiple excitation/emission events was accumulated on the ICCD sensor and read out at the end of the camera frame. The UV LED pulse width and pulse period, the intensifier delay time and on-time, the camera frame length (66.67 ms – 2 s) and the intensifier gain voltage could be varied independently. The source/camera timing parameters were the same for all of the time-resolved images and data presented here: excitation pulse width, 1500 μs, pulse period, 3000 μs, delay time, 10 μs, intensifier on-time, 1480 μs. All data reported here was acquired at a gain of 833 V. The camera control software enabled summation of multiple frames to yield a single composite. TIFF image with a bit depth equal to 1024 multiplied by the number of frames. All images reported here were summations of four frames (bit depth, 4096), and a feature of the camera control software was enabled that removes large variations in signal resulting from ion-feedback noise of the intensifier.

### Image Processing

Raw images were imported into NIH ImageJ (v1.42q) for all processing operations including cropping, contrast adjustment, and quantitative analysis.^*45*^ For each channel, 20 dark frames and 20 bright field images were stacked, converted to 32 bits, and median-filtered (radius 1), and each stack was averaged. The flat-field average was divided by the mean intensity of its central nine pixels to generate a normalized flat-field image. For each sample image, a median filter (radius 1) was applied and the master dark frame was subtracted. The resulting image was then divided by the normalized, master flat-field image, and the mean value of the detector offset was added back to the image. For ratiometric images and measurements, a binary mask was created by first averaging a series of Ypet images and then applying a threshold to highlight only regions exhibiting signal. The mask was applied to background-subtracted the FRET images were then divided by the donor image. Intensity-modulated ratiometric displays were generated using the Fire lookup table in ImageJ and a color lookup table was applied.

## ACKNOWLEDGMENTS

We thank Lumiphore, Inc. for providing Lumi4-Tb^®^. Funding was provided by NIGMS (R01 GM081030).

